# Modeling the Brain as a Shannon Information Source for fMRI-Based Network Analysis in Early Alzheimer’s Disease Diagnosis

**DOI:** 10.64898/2026.04.21.719889

**Authors:** Ulas Sedat Aydın, Abdulla Ahmadkhan, Gonul Gunal Degirmendereli, Fatos T. Yarman Vural

## Abstract

Alzheimer’s disease (AD) is an irreversible neurodegenerative disorder that gradually impairs memory, cognition, and behavior, making early diagnosis essential for slowing disease progression and improving patients’ quality of life. Functional Magnetic Resonance Imaging (fMRI) provides a noninvasive tool to study brain activity, yet many existing diagnostic models rely on black-box architectures that lack interpretability. In this study, we introduce a computational framework that models each anatomical brain region as a Shannon information source, thereby quantifying both the intrinsic information content of regions and the interactions among them. We used kernel density estimation to compute the probability density functions (PDFs) of voxel-level BOLD time series. From these PDFs, we derived regional entropy and pairwise Kullback-Leibler (KL) divergence measures. These measures were used to construct feature spaces representing information dynamics across the brain. We applied the framework to the ADNI resting-state fMRI dataset, which includes cognitively normal (CN), early mild cognitive impairment (EMCI), late mild cognitive impairment (LMCI), and AD subjects. Our findings indicate that entropy values increase with disease progression, while KL-based connectivity networks reveal a progressive loss of inter-regional interactions, especially in frontal, temporal, and parietal lobes. For classification, we trained multilayer perceptrons using voxel BOLD signals, entropy vectors, and KL divergence vectors. Models trained on KL features achieved the highest performance, outperforming both entropy-based and voxel-based approaches. These results demonstrate that the Shannon information source model offers an interpretable and statistically grounded approach for characterizing brain dynamics, while achieving superior diagnostic accuracy. Beyond AD, the proposed framework provides a generalizable tool for studying brain network alterations in neurological and psychiatric disorders.

## 1 Introduction

Alzheimer’s disease (AD) is the most common cause of dementia, accounting for 60–80% of cases worldwide. It is an irreversible and progressive neurodegenerative disorder that impairs memory, reasoning, and behavior, ultimately leading to loss of independence and death. According to the World Alzheimer Report 2024, more than 55 million people live with dementia worldwide and the global economic burden remains enormous (e.g., costs exceeded USD 1.3 trillion in 2018), underlining the public-health urgency of early, interpretable diagnostic biomarkers [1]. Although no definitive cure exists, early detection of AD is critical, as it enables timely interventions that can alleviate symptoms, slow disease progression, and improve patients’ quality of life.

Over the past two decades, advances in neuroimaging have enabled non-invasive exploration of the structural and functional alterations in the brain associated with AD. Functional Magnetic Resonance Imaging (fMRI) allows researchers to investigate patterns of brain connectivity and functional reorganization that occur during the early and clinical stages of the disease. Resting-state fMRI, in particular, captures dynamic neural activity and connectivity patterns that may reflect subtle functional disruptions prior to overt structural decline. When combined with machine learning (ML) and deep learning (DL) approaches, fMRI provides a promising avenue for developing sensitive biomarkers.

A wide range of ML and DL techniques have been applied to fMRI data for AD diagnosis, including Support Vector Machines (SVMs), Convolutional Neural Networks (CNNs), and Recurrent Neural Networks (RNNs) [2–9]. While these approaches report encouraging results, they are often criticized for relying heavily on voxel-level features and for functioning as “black-box” models that offer little interpretability regarding the underlying brain mechanisms. Such strategies may overlook the fundamental principles of information processing and flow within the brain, which are central to understanding neurodegeneration.

An alternative, principled perspective is provided by information theory, which leverages concepts such as entropy and divergence to quantify brain activity and connectivity [10, 11]. This approach is appealing because it provides interpretable, mathematically grounded measures of neural information processing. Shannon entropy measures the complexity or unpredictability of a signal, while Kullback–Leibler (KL) divergence quantifies the dissimilarity between probability distributions, thereby characterizing inter-regional information transfer. While these metrics have been applied to electroencephalography (EEG) and positron emission tomography (PET) studies for AD detection [11, 12], their application to region-wise fMRI modeling for AD remains relatively underexplored

Our work addresses this gap by integrating voxel-level probability modeling with region-wise information theory to construct effective feature representations for machine learning classification. We propose a computational framework for modeling the human brain as a network of Shannon information sources, where each anatomical region is treated as an information generator [13, 14]. From voxel-level fMRI signals, we estimate probability density functions (PDFs), compute Shannon entropy of each region, and measure KL divergence between all region pairs. Entropy values capture the information content of individual regions, while KL divergences quantify inter-regional information transfer. By constructing feature spaces from these measures, we investigate how information dynamics change across different stages of AD and evaluate their utility for early diagnosis.

We evaluate our approach using resting-state fMRI data from the Alzheimer’s Disease Neuroimaging Initiative (ADNI) and demonstrate that entropy- and KL-based models significantly outperform those using raw BOLD signals. These results highlight the potential of the proposed framework for interpretable and early-stage AD diagnosis. The key contributions of this study are as follows:

1. We introduce a Shannon information source model for brain regions that enables interpretable characterization of fMRI signals in terms of entropy and KL divergence.
2. We construct brain networks based on KL divergence, revealing how inter-regional information transfer deteriorates with AD progression.
3. We demonstrate that entropy- and KL-based features significantly outperform voxel-based features in classification with multilayer perceptrons.

The remainder of this paper is organized as follows. Section 2 describes the dataset and preprocessing pipeline, and presents the proposed information-theoretic model of the brain. Section 3 reports experimental results, including entropy analysis, KL-based brain networks, and classification performance. Section 4 discusses the findings in relation to existing literature and outlines directions for future research. Section 5 concludes the paper.

## 2 Materials and Methods

We used resting-state fMRI data from the Alzheimer’s Disease Neuroimaging Initiative (ADNI), a widely used benchmark dataset in neuroimaging-based Alzheimer’s research [2, 3]. Standard preprocessing pipelines were applied, including motion correction, slice timing correction, normalization, and spatial smoothing [7]. To model each anatomical brain region as an information source, we estimated probability density functions (PDFs) of voxel-level time series using Gaussian Kernel Density Estimation (KDE) [15]. From these PDFs, we computed Shannon entropy for each region and Kullback–Leibler (KL) divergence between all region pairs, capturing both local information complexity and inter-regional information flow.

Entropy and KL divergence values were then used to construct feature vectors for classification. For evaluation, multilayer perceptrons (MLPs) were trained using voxel-level signals, entropy features, and KL divergence features. Classification performance was compared across these three feature spaces to assess the added value of information-theoretic modeling.

### 2.1 Dataset

We evaluated our approach using resting-state functional Magnetic Resonance Imaging (rs-fMRI) data obtained from the Alzheimer’s Disease Neuroimaging Initiative (ADNI). The dataset included subjects from four clinical groups: cognitively normal (CN), early mild cognitive impairment (EMCI), late mild cognitive impairment (LMCI), and Alzheimer’s disease (AD). To avoid potential biases in the subsequent classification tasks, we selected a balanced subset of subjects across the four groups. In total, 31 subjects were included, with 20 sessions per subject and 140 time samples per session, resulting in a total of 11,200 fMRI data instances (subjects × sessions). The detailed distribution is provided in Table 1.

**Table 1:**
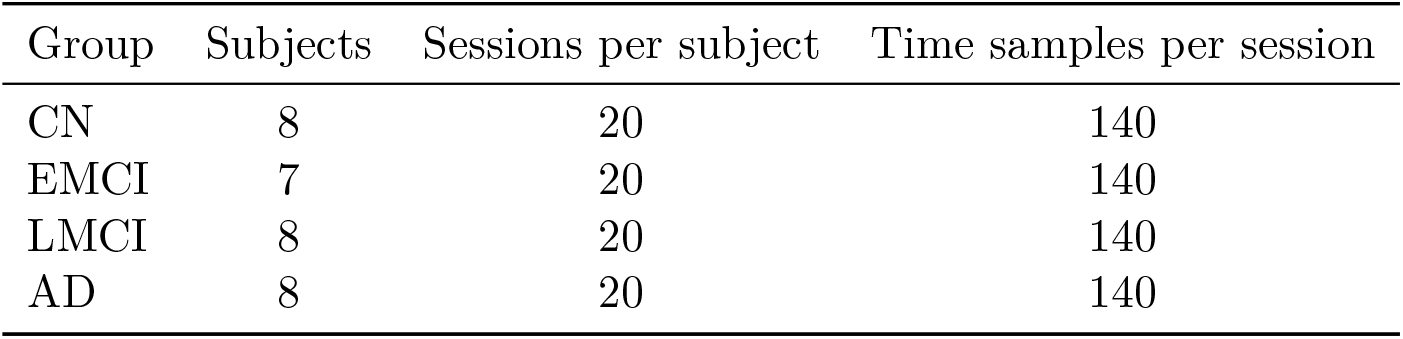
Subject distribution in the ADNI-fMRI dataset.

### 2.2 Preprocessing

All fMRI data were preprocessed using the Statistical Parametric Mapping (SPM) software package. The pipeline consisted of the following steps:

1. Slice timing correction– to correct temporal shifts introduced during sequential slice acquisition.
2. Motion correction (realignment) – to reduce artifacts caused by head movement across scans.
3. Normalization – spatially warping images into the standard MNI space to allow group-level comparisons.
4. Smoothing – applying a Gaussian kernel to reduce noise and increase signal-to-noise ratio.
5. Parcellation – mapping voxels to anatomical regions using the Automated Anatomical Labeling (AAL) atlas. The first 90 regions of the AAL atlas were used, as they are highly relevant to AD-related alterations.

This preprocessing ensured that each voxel’s BOLD time series could be reliably assigned to a specific anatomical region.

### 2.3 Information-Theoretic Modeling of Brain Regions

We model each anatomical region as a Shannon information source. The central idea is that the BOLD time series within a region represent stochastic observations generated by an underlying neural process.

#### 2.3.1 Entropy Estimation

The entropy of a region reflects the amount of information encoded by its voxel activity. Structured and coherent patterns in active regions are expected to yield lower entropy, while random fluctuations in less active or disrupted regions yield higher entropy. [16]. To estimate the entropy of each anatomical region, we treat voxel-level BOLD responses as realizations of a random variable and apply kernel density estimation (KDE) to model their probability distribution.

Let *v*_*i*_(*t*) denote the voxel intensity at time *t* for voxel *i* within region *s*. We estimate the probability density function (PDF) via Gaussian kernel density estimation (KDE):

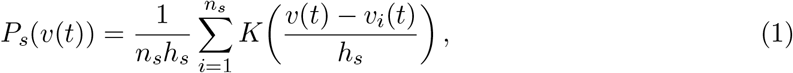

where *n*_*s*_ is the number of voxels in region *s, h*_*s*_ is the bandwidth, and *K*(·) is a Gaussian kernel. The Shannon entropy for region *s* is then computed (in discrete form) as:

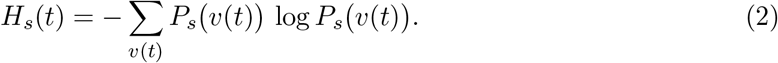

#### 2.3.2 KL Divergence Estimation

The representation of the brain regions as information sources facilitates the estimation of their connectivity with KL divergence. The KL divergence between regions quantifies the statistical dissimilarity between their PDFs, thereby measuring the extent of information transfer. Thus, KL divergence values between anatomical region pairs are used as the connection weights of the brain network formed by these information sources. [15].

To estimate the brain networks, we utilize voxel density values *v*_*i*_(*t*) measured at location (*x*_*i*_, *y*_*i*_, *z*_*i*_) and time *t*. For every information source **s** and time sample *t*, the probability density function *P*_*s*_(*v*(*t*)) for all voxels in the source is estimated using the mentioned equation (1). KL divergence between two anatomical regions *k* and *l* for ∀ *k*≠ *l* at time *t* is estimated as:

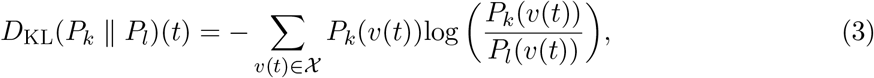

Such that 𝒳 is the set of voxel values at time *t* in either the anatomical region *k* or *l*.

A smaller KL value indicates stronger similarity (and thus stronger interaction) between regions. Using this measure across all pairs of regions, we construct weighted brain networks that capture connectivity patterns at each disease stage.

### 2.4 Feature Construction

Two types of feature vectors were derived:

1. Entropy vectors (90-dimensional): Each dimension corresponds to the entropy of one anatomical region.
2. KL divergence vectors (90 × 90 = 8100-dimensional): Each entry corresponds to the KL divergence between a pair of regions.

These features provide a compact and interpretable representation of brain activity and connectivity.

### 2.5 Classification with Multilayer Perceptrons

To evaluate the discriminative power of the proposed features, we trained multilayer perceptrons (MLPs) on three feature sets to classify four stages of the Alzheimer’s disease: (1) averaged BOLD signals, (2) entropy vectors, and (3) KL divergence vectors.

We chose MLPs—one of the simplest classifier types—primarily to emphasize that the performance difference stems from the feature space rather than model complexity.

The network architecture consisted of three fully connected layers with ReLU activations and a final softmax output layer for multi-class classification. The dataset was split into training (90%) and testing (10%) using stratified sampling, and performance was evaluated using 10-fold cross-validation.

Model performance was assessed using Precision, Recall, F1-score, and Accuracy.

## 3 Results

### 3.1 Entropy Analysis

We first compared the entropy values estimated for the 90 anatomical regions across groups. Entropy values were lowest in the cognitively normal (CN) group, higher in early mild cognitive impairment (EMCI), and highest in AD patients, indicating increasing disorder in voxel-level activity with disease progression. See Figure 1, which illustrates mean entropy values for a selection of 50 regions characterized by the lowest entropy in healthy controls.

**Figure 1:**
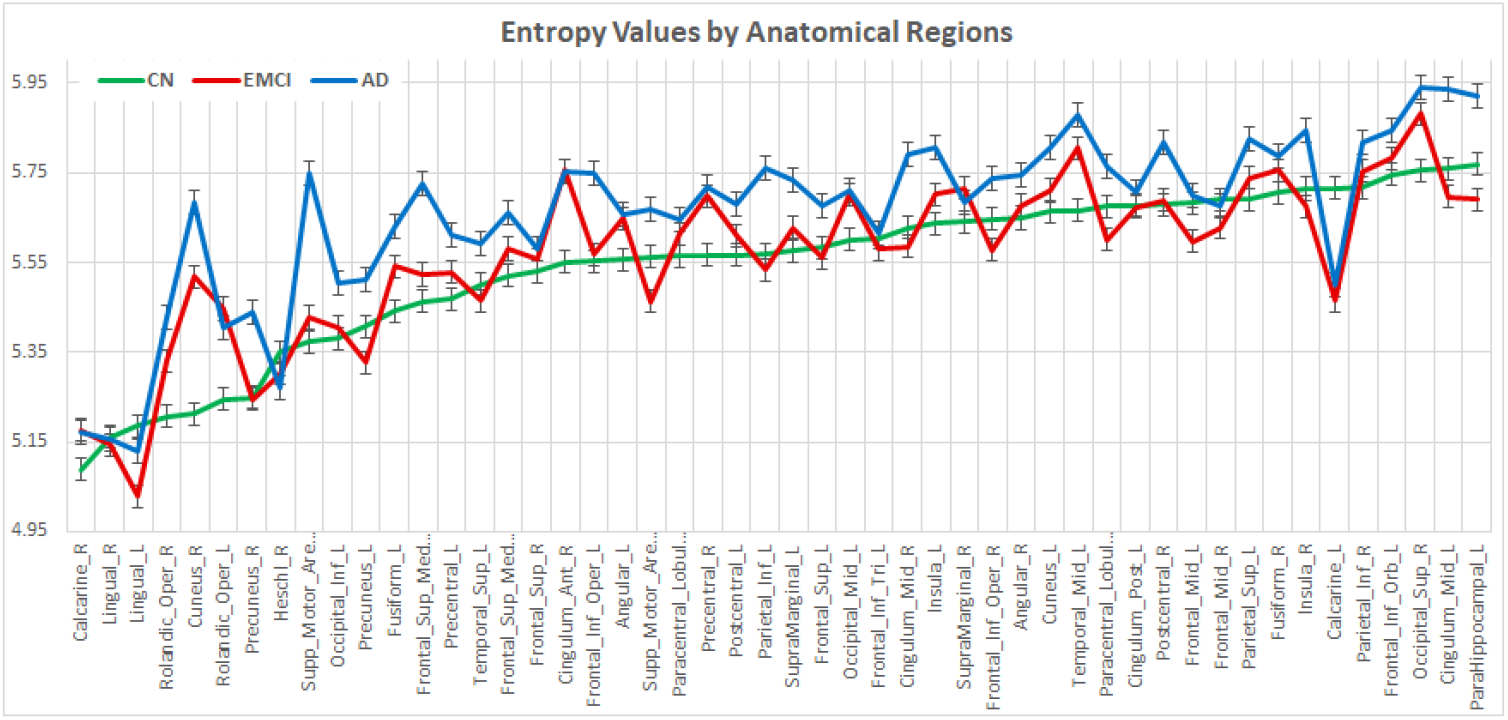
Entropy values across diagnostic groups. Entropy significantly increased from CN to AD, indicating higher irregularity in BOLD signals. Error bars represent standard error.

### 3.2 KL Divergence Brain Networks Analysis

To further investigate inter-regional information transfer, KL divergence values were computed between all pairs of anatomical regions. We generated arc-weight matrices, where KL divergence measures represent the degree of co-activation between pairs of the 90 anatomical regions. Therefore, each matrix has 90×90 = 8100 connections for each time instant. Group-wise KL divergence matrices were thresholded to highlight the strongest connections, allowing the construction of functional brain networks. To identify the most robust connections, we averaged KL divergence matrices across all time instances and then thresholded the resulting group-average matrix to retain only the strongest (lowest divergence) connections.

The results revealed that CN subjects exhibited high inter-lobe connectivity, with strong links particularly in the default mode network (DMN) and between frontal and temporal lobes. In EMCI patients, connectivity began to weaken, especially in the temporal and parietal regions, though some hub structures remained intact. Finally, AD patients exhibited severe disintegration, characterized by sparse and fragmented networks with reduced long-range connectivity.

These findings indicate that KL divergence analysis captures progressive disruptions in brain information transfer, consistent with the hypothesis that AD is a disconnection syndrome. Notably, the reduction in cross-lobe connectivity provides a quantitative explanation for the cognitive impairments observed in memory, reasoning, and attention as the disease advances.

### 3.3 Visualization of Brain Networks

To provide an intuitive representation of these disruptions, we visualized the KL divergence networks for each group using BrainNet Viewer. Anatomical nodes were defined according to the AAL atlas. Node colors represent the entropy values of brain regions (blue = low entropy, red = high entropy). Edge visibility indicates edge strength; stronger connections are shown with more prominent lines, and weaker connections are shown with fainter lines.

Figures 2–4 illustrate the resulting networks for cognitively normal (CN), early mild cognitive impairment (EMCI), and Alzheimer’s disease (AD) groups.

**Figure 2:**
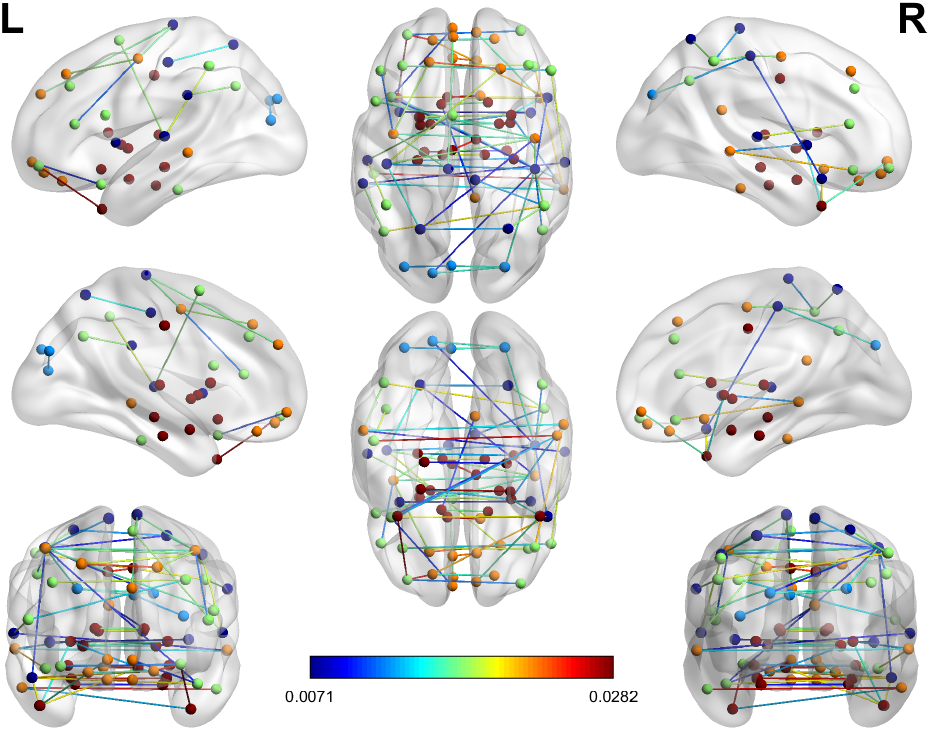
Estimated KL-based brain network for CN subjects.

**Figure 3:**
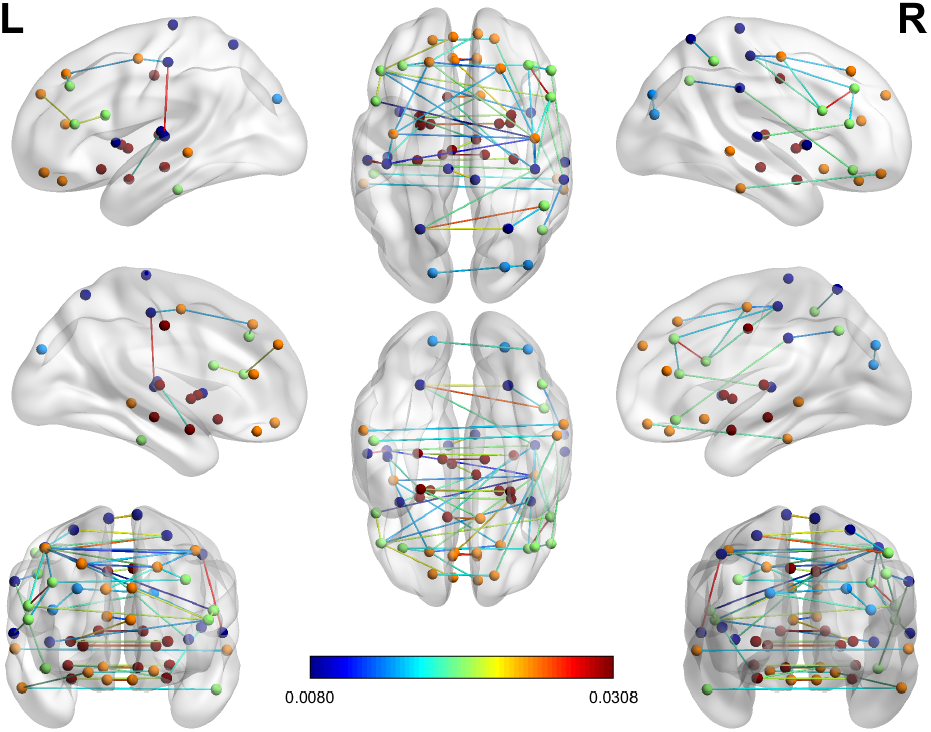
Estimated KL-based brain network for EMCI subjects.

**Figure 4:**
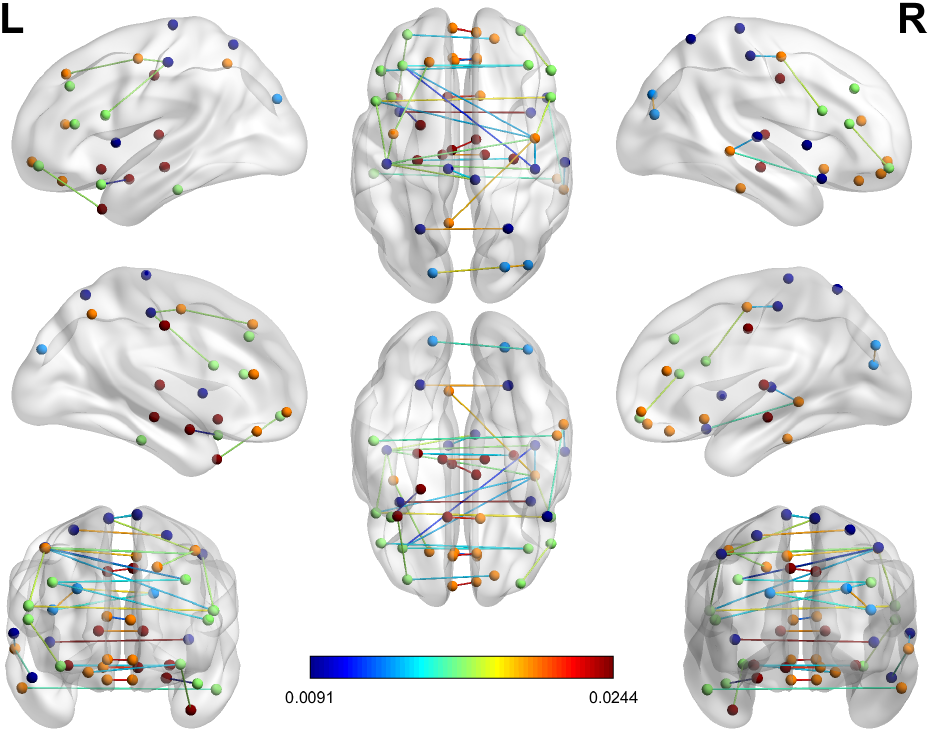
Estimated KL-based brain network for AD subjects.

Connectivity decreased markedly from CN to EMCI to AD. In the CN group, a well-organized network structure is observed, with strong interhemispheric and long-range connections linking posterior cingulate cortex, medial prefrontal cortex, and inferior parietal regions—key nodes of the default mode network (DMN). In the EMCI group, the network shows early signs of disconnection, particularly in long-range frontal–parietal and interhemispheric pathways, although the core DMN structure remains partially preserved. By contrast, the AD group exhibits a markedly fragmented network: long-range connections are largely disrupted, and connectivity is confined to a few localized clusters. These observations are consistent with the well-documented “disconnection syndrome” in AD, whereby progressive neurodegeneration leads to collapse of large-scale integration in the DMN while sparing some local connectivity.

These visualizations provide clear qualitative support for the quantitative results obtained with entropy and KL divergence, reinforcing the interpretation that AD progression is marked by both elevated local signal irregularity and loss of inter-regional information flow. Overall, these findings suggest that visualization of KL divergence–based brain networks can capture disease-related disruptions that may serve as potential imaging biomarkers for early and advanced stages of Alzheimer’s disease.

### 3.4 Classification Performance

We trained multilayer perceptrons (MLPs) with three feature sets: BOLD averages, entropy vectors, and KL divergence vectors. Performance metrics are summarized in Table 2. KL-based features achieved the highest accuracy (93.6%), followed by entropy features (82.8%), while BOLD-based features performed less effectively (73.6%). KL features also yielded the highest precision, recall, and F1 scores.

**Table 2:** MLP classification performance across feature types.

| Input Features | Precision | Recall | F1 | Accuracy |
| --- | --- | --- | --- | --- |
| BOLD | 0.791 | 0.736 | 0.740 | 73.6% |
| Entropy | 0.811 | 0.828 | 0.790 | 82.8% |
| KL | 0.931 | 0.914 | 0.911 | 93.6% |

These results demonstrate that information-theoretic features not only capture disease-related changes more effectively than raw voxel signals but also enhance the performance of relatively simple classifiers.

### 3.5 Results Summary

Our results indicate a consistent increase in entropy values with disease progression, reflecting greater signal irregularity in AD-affected brains. Analysis of KL divergence networks revealed progressive reductions in inter-regional information transfer, especially across parietal and temporal regions [11]; this observation is consistent with EEG and connectivity literature reporting decreased coherence and information transfer in frontal, temporal and parietal areas in AD [17]. Visualization of the brain networks further confirmed these patterns, showing a systematic loss of connectivity from CN to AD. These findings align with known patterns of connectivity disruption in AD [18].

Classification experiments demonstrated that KL-based feature vectors achieved the highest performance in our setting, surpassing entropy-based and voxel-based approaches. This result is consistent with literature suggesting that carefully designed, interpretable feature representations can match or improve performance compared with voxel-level black-box models in certain settings, while also providing greater mechanistic insight [19, 20]. It also align with our earlier information-theoretic modeling of fMRI [13, 15].

## 4 Discussion

Our findings demonstrate that modeling brain regions as Shannon information sources provides a principled and interpretable framework for characterizing functional alterations in Alzheimer’s disease. The observed entropy increases are consistent with prior EEG studies reporting higher signal complexity in AD patients [10], while reduced KL divergence aligns with PET-based findings of disrupted metabolic connectivity [11]. Beyond numerical analyses, the visualization of KL-based brain networks revealed a progressive sparsification of connectivity, particularly in frontal, temporal, and parietal lobes. Dense and symmetric networks observed in CN subjects gradually weakened in EMCI and LMCI, and by the AD stage only localized clusters with residual entropy survived. This convergence of quantitative and visual evidence strengthens the interpretation that AD progression is associated with both elevated local signal irregularity and disrupted inter-regional information transfer.

Compared to existing machine learning and deep learning approaches that rely on voxel-level features, our model provides a principled and interpretable alternative. While black-box models such as CNNs or RNNs have achieved high accuracy [2, 3, 8], their limited interpretability poses challenges for clinical adoption. By contrast, entropy and KL divergence features are mathematically grounded, biologically meaningful, and directly linked to information processing in neural systems. This transparency makes them promising candidates for biomarker development. Recent work by Vlontzou et al. (2025) further demonstrates that interpretable machine learning frameworks achieve robust and clinically meaningful AD diagnosis, reinforcing the value of transparency in neuroimaging-based models [20]. Similarly, Bloch and Friedrich (2022) showed that explainability methods such as SHAP can enhance black-box classifiers while maintaining competitive accuracy, further highlighting the importance of interpretability for clinical adoption [19].

The KL-based visualizations further enhance interpretability by making topological disruptions in brain networks tangible. Healthy subjects showed strong inter-lobe connectivity with balanced hubs, whereas AD subjects exhibited fragmented networks dominated by weakly connected subgraphs. These findings align with clinical knowledge that Alzheimer’s disease is characterized by large-scale network disconnection, prominently affecting hubs of the Default Mode Network (DMN) and their integration with fronto-parietal systems [18, 21]. Importantly, the visual evidence provides an intuitive bridge between quantitative connectivity metrics and their potential clinical interpretation.

Entropy increases observed across disease stages also align with the hypothesis that AD disrupts structured neuronal firing, leading to more random and less predictable activity patterns. This is consistent with previous EEG-based findings of higher complexity in AD patients [12]. In parallel, KL divergence captures the progressive decoupling of information exchange between regions, a hallmark of impaired synaptic communication and network integrity.

Despite these promising results, several limitations should be noted. The dataset size was moderate, and only resting-state fMRI was used. Moreover, classification relied on multilayer perceptrons; future work should test whether combining information-theoretic features with more advanced classifiers (e.g., graph neural networks) could further improve performance. Incorporating multimodal imaging data (e.g., PET, EEG) may also strengthen clinical applicability [3]. Finally, graph-theoretical analyses (e.g., node degree, clustering coefficient, eigen decomposition) of KL-based networks represent a natural next step to systematically quantify the topological alterations suggested by the visualizations.

In summary, the integration of entropy analysis, KL-based connectivity, and visualization provides a comprehensive framework for studying Alzheimer’s disease. The combined numerical and visual evidence supports the conclusion that AD progression is characterized by both increased local signal irregularity and reduced inter-regional information transfer. This dual perspective—quantitative and qualitative—positions the Shannon information source model as a powerful and interpretable tool for early diagnosis and clinical understanding of neurodegeneration.

## 5 Conclusion

In this study, we presented a novel information-theoretic framework for modeling the brain as a network of Shannon information sources. By representing each anatomical region as an information generator, we quantified both local information content via entropy and inter-regional information transfer via Kullback–Leibler (KL) divergence. Applied to resting-state fMRI data from the ADNI cohort, our approach revealed progressive increases in entropy and reductions in KL-based connectivity with Alzheimer’s disease (AD) progression. These alterations captured meaningful disruptions in brain dynamics that were not apparent from raw BOLD signals.

Classification experiments demonstrated that entropy- and KL-based features significantly outperformed voxel-level BOLD features, with KL features achieving the highest accuracy of 93.6% using a multilayer perceptron. Beyond their predictive performance, the proposed measures are interpretable and mathematically grounded, offering insights into disease-related changes in neural information processing.

This work highlights the promise of information-theoretic approaches for early detection of AD. By bridging computational neuroscience and clinical neuroimaging, the Shannon information source model offers both practical diagnostic tools and theoretical insight into neurodegeneration. Future studies with larger datasets, multimodal imaging, and advanced classifiers may further validate and extend the utility of this framework to other neurological and psychiatric disorders.

## Declarations

## Acknowledgments

We would like to thank Dr. Fırat Soylu for his valuable contributions in the preprocessing phase, Prof. Hüseyin Boyacı for his knowledge and experience on fMRI, and Abdul Sami Asif for his valuable support during the training process of the model.

## Author Contributions

FYV: Conceptualization, Formal Analysis, Methodology, Project Administration, Supervision, Validation, Writing – Review & Editing. GGD: Conceptualization, Data Curation, Formal Analysis, Methodology, Software, Supervision, Visualization, Writing – Original Draft. USA: Data Curation, Formal Analysis, Investigation, Software, Visualization, Writing – Original Draft. AA: Formal Analysis, Software, Visualization, Writing – Original Draft.

## Data Availability

The Image and Data Archive (IDA): https://ida.loni.usc.edu

